# Root meristem growth factor 1 controls root meristem size through reactive oxygen species signaling

**DOI:** 10.1101/244947

**Authors:** Masashi Yamada, Xinwei Han, Philip. N. Benfey

**Affiliations:** Department of Biology and HHMI, Duke University, Durham, NC 27710, USA

## Abstract

Stem cell niche and root meristem size are maintained by intercellular interactions and signaling networks of a plant peptide hormone, Root Meristem Growth Factor 1 (RGF1). How RGF1 regulates root meristem development is an essential question to understand stem cell function. Although five receptors of RGF1 have recently been identified, the downstream signaling mechanism remains unknown. Here, we report a series of signaling events following RGF1 action. The RGF1-receptor pathway controls distribution of reactive oxygen species (ROS) along the developmental zones of the *Arabidopsis* root. We identify a novel transcription factor, *RGF1 INDUCIBLE TRANSCRIPTION FACTOR 1* (*RITF1*), which plays a central role in mediating RGF1 signaling. Manipulating *RITF1* expression leads to redistribution of ROS along the root developmental zones. Changes in ROS distribution, in turn, enhance the stability of the PLETHORA2 (PLT2) protein, a master regulator of root stem cells. Taken together, our study clearly depicts a signaling cascade initiated by RGF1 and links the RGF1 peptide to ROS regulatory mechanisms.

## Text

Roots encounter various environmental conditions in soil and respond by altering their growth. The *Arabidopsis* root has a simple cylindrical structure. Each layer of the cylinder consists of a different cell-type, which is generated from a stem cell population at the root tip, and develops into mature cells along the longitudinal axis of the root. Developmental stages are defined as the meristematic zone, the elongation zone, and the differentiation zone. Root growth arises through controlled cell division in the meristematic zone and subsequent cell elongation and differentiation in the elongation and differentiation zones. During the transition to differentiation, most cells arrest division and increase their size through post-mitotic cell expansion. The size of the developmental zones is determined by intrinsic and extrinsic signals. In the *Arabidopsis* root, superoxide (O_2_^-^) and hydrogen peroxide (H_2_O_2_) exhibit distinct distribution patterns along the developmental zones^1^. Superoxide primarily accumulates in dividing cells in the meristematic zone, while hydrogen peroxide mainly accumulates in elongated cells in the differentiation zone^1,2^. The balance between O_2_^-^ and H_2_O_2_ modulates the transition from proliferation to differentiation^2^. The *UPBEAT1* (*UPB1*) gene regulates meristematic zone size by restricting H_2_O_2_ distribution in the elongation zone^2^. These findings have demonstrated that reactive oxygen species (ROS) are an intrinsic signal involved in establishing the size of the meristematic zone.

The RGF1 peptide is also able to control the size of the meristematic zone both as an intrinsic signal and when extrinsically applied^3–5^. External treatment with RGF1 increases the size of the meristematic zone, while the *rgf1/2/3* triple mutant has a smaller meristematic zone^3^. Five receptor-like kinases have been identified as RGF1 receptors^6–8^. Quintuple mutants of these receptors lack most of the cells in the root meristem and are insensitive to RGF1, demonstrating that the RGF signaling pathway controls root meristem size via these receptors^8^. The RGF1 signaling pathway controls the stability of the PLETHORA (PLT) 1/2 proteins^3^, which are required for stem cell maintenance^9^. However, it is not known how RGF modulates the size of the meristematic zone and the stability of the PLT1/2 proteins. Here, we show that the RGF1 signaling pathway modulates ROS distribution along the three root developmental zones effectively controlling the size of the meristematic zone. Changes in ROS distribution result in changes in stability of the PLT2 protein. Transcriptome analysis after RGF1 treatment identified elevated expression of a meristematic zone-specific novel transcription factor (*RITF1*). Over-expression of the *RITF1* gene phenocopies the enlarged meristematic zone and altered distribution of ROS signaling upon RGF1 treatment.

It has been reported that RGF1 modulates meristematic zone size^3,6–8^. To confirm the effect of RGF1 treatment on the meristematic zone with greater specificity, we used the meristematic zone-specific marker HIGH PLOIDY2 (HPY2)-GFP protein^10^ (Fig. 1a and b). HPY2-GFP was detected in an enlarged area 24h after RGF1 treatment (Fig. 1a-e) and this correlated with a larger meristematic zone (Fig. 1c and d). These results combined with defects in the meristematic zone in *rgf* and *rgf1 receptor* (*rgfr*) mutants suggest that RGF1 controls gene expression primarily in the meristematic zone. Therefore, we performed a meristematic zone-specific transcriptome analysis to uncover the molecular mechanism underlying the RGF1 signaling pathway. To identify primary target genes upon RGF1 treatment, we isolated the meristematic zone based on the *HPY2-GFP* signal one hour after RGF1 treatment (Fig. 1f).

**Figure 1.**
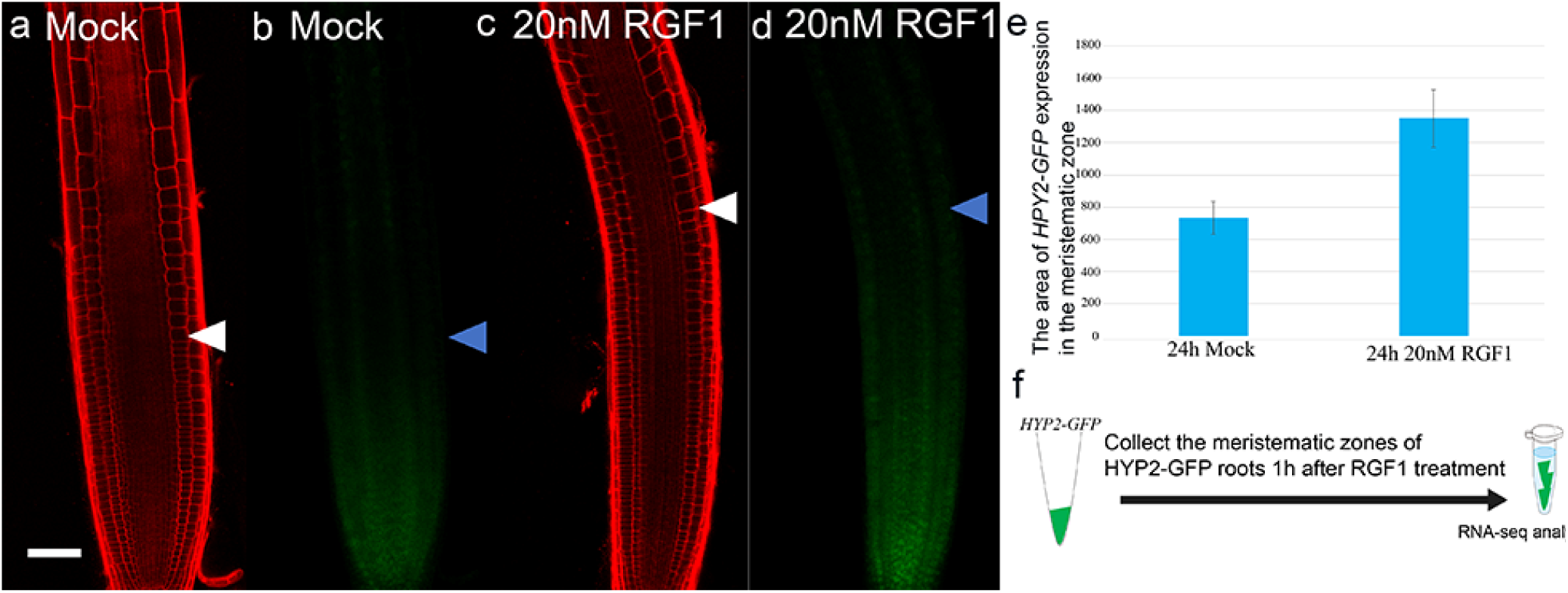
Expression of meristematic zone marker and transcriptome analysis upon RGF1 treatment. Confocal images of *HPY2-GFP* are shown 24 h after treatments with water (mock) (a and b) and 20 nM RGF1 (c and d). Seedlings were grown on MS medium for seven days before treatment. Propidium iodide stained roots (a and c); GFP signals (b and d). White and blue arrow heads indicate the junction between the meristematic zone and the elongation zone. Scale bar = 50 µm (e) Area (µm^2^) of *HPY2-GFP* expression is shown. Error bars represent ± standard deviation (SD; n≥9). (f) Schematic of RNA extraction following RGF1 treatment.

Since the expression of *HPY2-GFP* and the size of the meristematic zone were unchanged in this time period (data not shown), we can exclude the possibility that an enlarged meristem is the reason for elevated RNA levels. Expression of 583 genes was significantly altered in the RNA-seq data between RGF1 and mock treatment (FDR-adjusted p value < 0.1) (Extended Data). Most differentially expressed genes were positively regulated by RGF1 treatment. However, our transcriptome analysis revealed specific down-regulation of *RGF1* itself, suggesting negative feed-back regulation (Extended Data). Significantly enriched Gene Ontology (GO) categories included “oxidoreductase activity” (p= 4.90E-06) and “oxidation reduction” (p= 4.90E-05) (Extended Data Fig. 1 and Extended Data). These data suggested that RGF1 might signal through a ROS intermediate to control the size of the meristematic zone.

To examine the relationship between RGF1 and ROS signaling, we analyzed the distribution of superoxide and hydrogen peroxide after RGF1 treatment. The specific indicator H_2_O_2_-3’-O-Acetyl-6’-O-pentafluorobenzenesulfonyl-2’-7’-difluorofluorescein-Ac (H_2_O_2_-BES-Ac)^2^ for hydrogen peroxide exhibited lower fluorescence in the meristematic and elongation zones 24 h after RGF1 treatment (Fig. 2a and c). Superoxide signals were detected by nitroblue tetrazolium (NBT) staining^1^ and observed more broadly in the meristematic and elongation zone 24 h after RGF1 treatment (Fig. 2b and d). To determine if these changes in ROS distributions depend on the RGF1 receptors, ROS signals were detected in the *rgfr1/2/3* triple mutant. The *rgfr1/2/3* triple mutant was insensitive to RGF1 and did not form a larger meristematic zone upon RGF1 treatment (Fig. 2e). Levels of H_2_O_2_ and O_2_^-^ in the *rgfr1/2/3* triple mutant were comparable between mock and RGF1 treatments (Fig. 2e-h). These data are consistent with our hypothesis that the RGF1-receptor signaling pathway controls the size of the meristematic zone via ROS.

**Figure 2.**
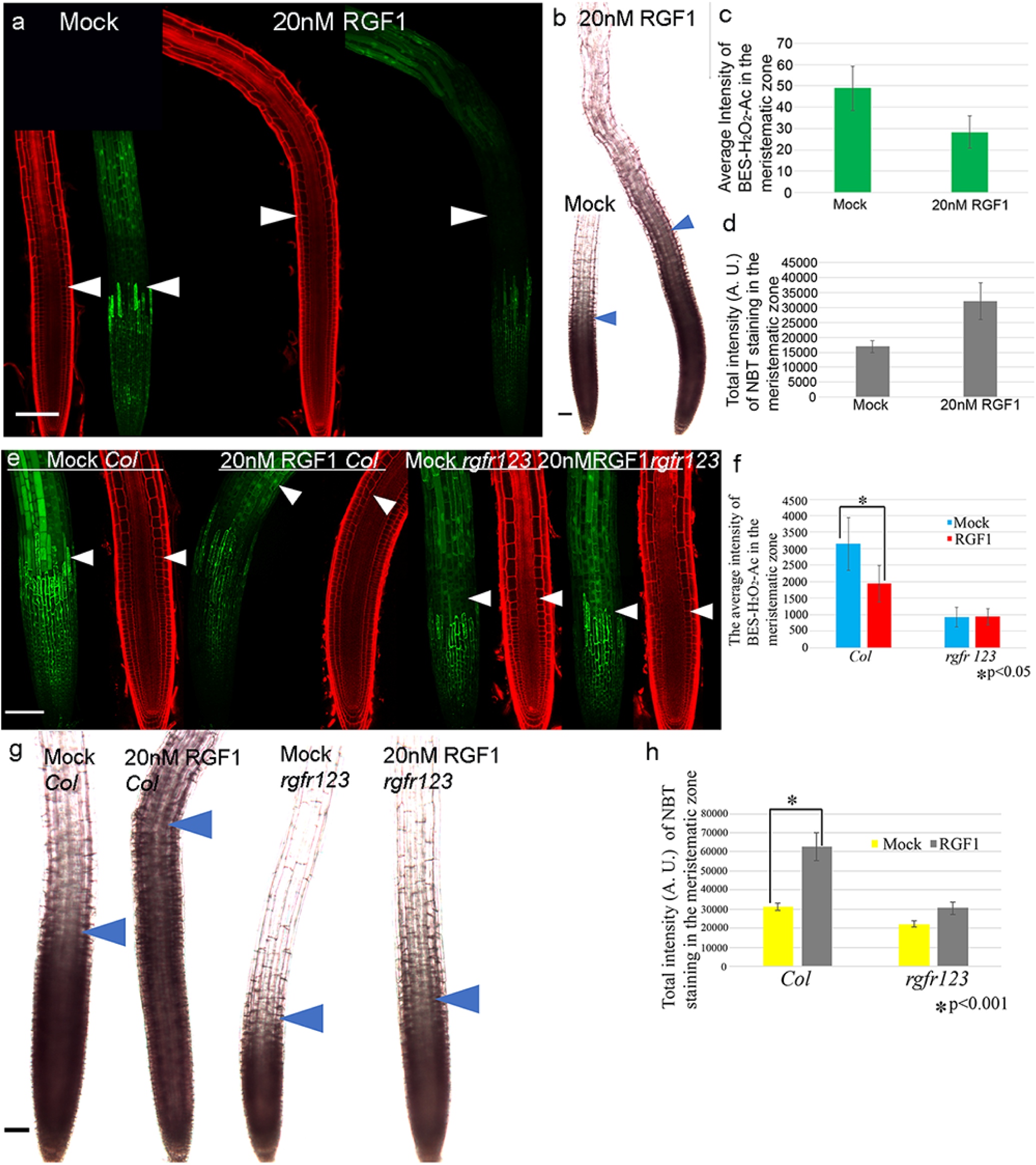
Distribution of ROS levels upon RGF1 treatment. (a). Confocal images of roots 24 h after treatment with Water (mock) and 20 nM RGF1 (right). Propidium iodide (PI) staining (red images), H_2_O_2_-BES-Ac fluorescence (green images). (b) Roots stained with NBT 24 h after treatment with mock or 20 nM RGF1 (right). (c) Quantification of H_2_O_2_-BES-Ac intensity in meristematic zone (n = 6, ± SD; p < 0.003). (d) Quantification of NBT staining intensity (arbitrary units (A.U.) values) in meristematic zone (n≥8, ± SD; p < 0.001). (e) Confocal images of roots 24 h after treatment with mock or 20 nM RGF1 in wild type (Col-0) (left four images) or the *rgfr 1/2/3* triple mutant (right four images). PI staining (red images). H_2_O_2_-BES-Ac fluorescence (green images). (f) Quantification of H_2_O_2_-BES-Ac staining intensity in the meristematic zone in wild type and the *rgfr 1/2/3* triple mutant (n = 5, ± SD; *p < 0.05). (g) Roots stained with NBT 24 h after treatment with mock or 20 nM RGF1 in wild type (left two images) or the *rgfr 1/2/3* mutant (right two images). (h) Quantification of NBT staining intensity (A.U. values) in the meristematic zone in wild type or the *rgfr 1/2/3* triple mutant (n = 5, ± SD; *p < 0.001). White and blue arrowheads indicate the junction between the meristematic zone and elongation zone. Scale bar = 50 µm. Seedlings were grown on MS medium for seven days before treatment.

It was previously reported that the RGF1 signaling pathway regulates PLT1/2 post-translationally^6^. We compared signals from a *PLT2* transcriptional fusion line; *promoter PLT2* (*pPLT2*)-*CFP^11^* and from a *PLT2* translational fusion line; *genomic PLT2* (*gPLT2*)-*YFP^11^* (Extended Data Fig. 2a-c) and observed broader localization of *gPLT2*-*YFP* signals 24 h after RGF1 treatment (Extended Data Fig. 2b-d). However, the levels and localization of *pPLT2*-*CFP* were comparable between Mock and RGF1 treatments (Extended Data Fig. 2a), even though RGF1-treated roots had a larger meristematic zone. These experiments confirmed that RGF1 regulates PLT2 post-translationally. We detected the highest levels of *gPLT2*-*YFP* and *pPLT2*-*CFP* in the cells around the QC (Extended Data Fig. 2a and b). The *gPLT2*-*YFP* signal gradually declined in the meristem after mock treatment (Extended Data Fig. 2b). However, after RGF1 treatment, the *gPLT2*-*YFP* signal decreased more gradually and was broadly localized in a larger meristematic zone (Extended Data Fig. 2c). Since proteasome-dependent degradation of proteins occurs in the presence of elevated H_2_O_2_ levels^12,13^, we hypothesized that broader localization of the PLT2 protein is due to the decreased H_2_O_2_ and increased O_2_^-^ levels upon RGF1 treatment as shown in Figure 2. To determine if H_2_O_2_ can decrease the stability of the PLT2 protein, we treated the *gPLT2*-*YFP* line with RGF1 and H_2_O_2_. H_2_O_2_ treatment inhibited the size increase of the meristem upon RGF1 treatment (Extended Data Fig. 3d and e). Furthermore, *gPLT2*-*YFP* signals were not localized as broadly in roots co-treated with RGF1 and H_2_O_2_ as compared with roots only treated with RGF1 (Extended Data Fig. 3b and c). These results are consistent with our hypothesis that lower H_2_O_2_ levels enhance the stability of the PLT2 protein. To further probe the relationship between PLT2 protein stability and ROS, we measured *gPLT2*-*YFP*, O_2_^-^, and H_2_O_2_ in a shorter time course (4-10h) after RGF1 treatment. At 6 h after RGF1 treatment, we began to detect broader localization of *gPLT2*-*YFP* (Fig. 3a and b, Extended Data Fig. 4). At the same time, increased superoxide levels, as indicated by broader staining with NBT, were observed in the meristematic zone (Fig. 3g and h) along with lower signals of H_2_O_2_-BES-Ac in the distal side of the meristematic zone (Fig. 3m and n, white arrows). At 8 h and 10 h after RGF1 treatment, expanded *gPLT2*-*YFP* expression and O_2_^-^ signals correlated with declining H_2_O_2_ signals (Fig. 3c-f, 3i-l, 3o-r and Extended Data Fig. 4a-c). These data indicate that RGF1 treatment increases root meristem size and PLT2 protein stability by modulating ROS levels.

**Figure 3.**
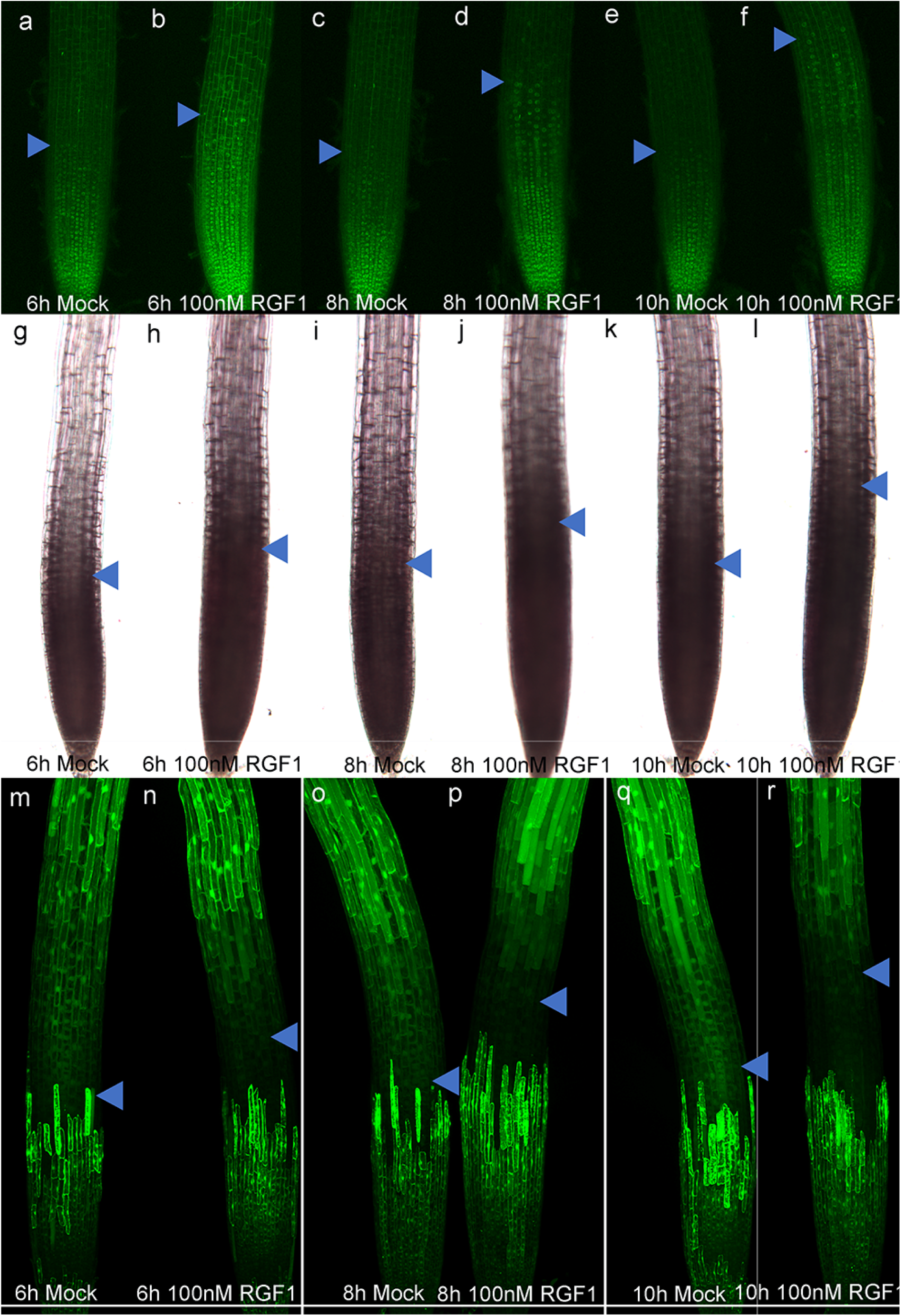
Localization of gPLT2-YFP, NBT, and H_2_O_2_-BES-Ac after RGF1 treatment. Localization of gPLT2-YFP, NBT, (M-R) H_2_O_2_-BES-Ac, 6 h after treatment with water (mock) (a, g, and m) or 100 nM RGF1 (b, h, and n), 8 h after treatment with mock (c, i, and o), or 100 nM RGF1 (d, j, and p). 10h after treatment with mock (e, k, and q) or 100nM RGF1 (f, l, and r). Blue arrow heads indicate junction between meristematic zone and elongation zone. Scale bar = 50 µm. Seedlings were grown on MS agar plates for seven days before treatment.

To identify for downstream factors that mediate the RGF1/ROS signaling pathway, we combined our RGF1 treatment transcriptome data with our previously published transcriptome data from the three root developmental zones^14^. We looked for genes whose expression was induced by RGF1 and specific to the meristematic zone. The *PLANT AT-RICH SEQUENCE and ZINC-BINDING TRANSCRIPTION FACTOR* (*PLATZ*) *FAMILY PROTEIN* gene (AT2G12646) is strongly expressed in the meristematic zone with lower expression in the elongation zone and greatly reduced expression in the differentiation zone (Extended Data Fig. 5a). As this gene has not been characterized, we named it *RGF1 INDUCIBLE TRANSCRIPTION FACTOR* 1 (*RITF1*). Expression of *RITF1* increased approximately 2-fold after 1 h of RGF1 treatment (Extended Data and Extended Data Fig. 5b). To understand its function, *RITF1*was inducibly over-expressed using the estradiol inducible promoter system^15^. After 24 h of 10µM β-estradiol treatment, the meristematic zone became enlarged and the number of cells increased as compared with mock treatment (Fig. 4a and b). These phenotypes are very similar to RGF1 treated roots.

**Figure 4.**
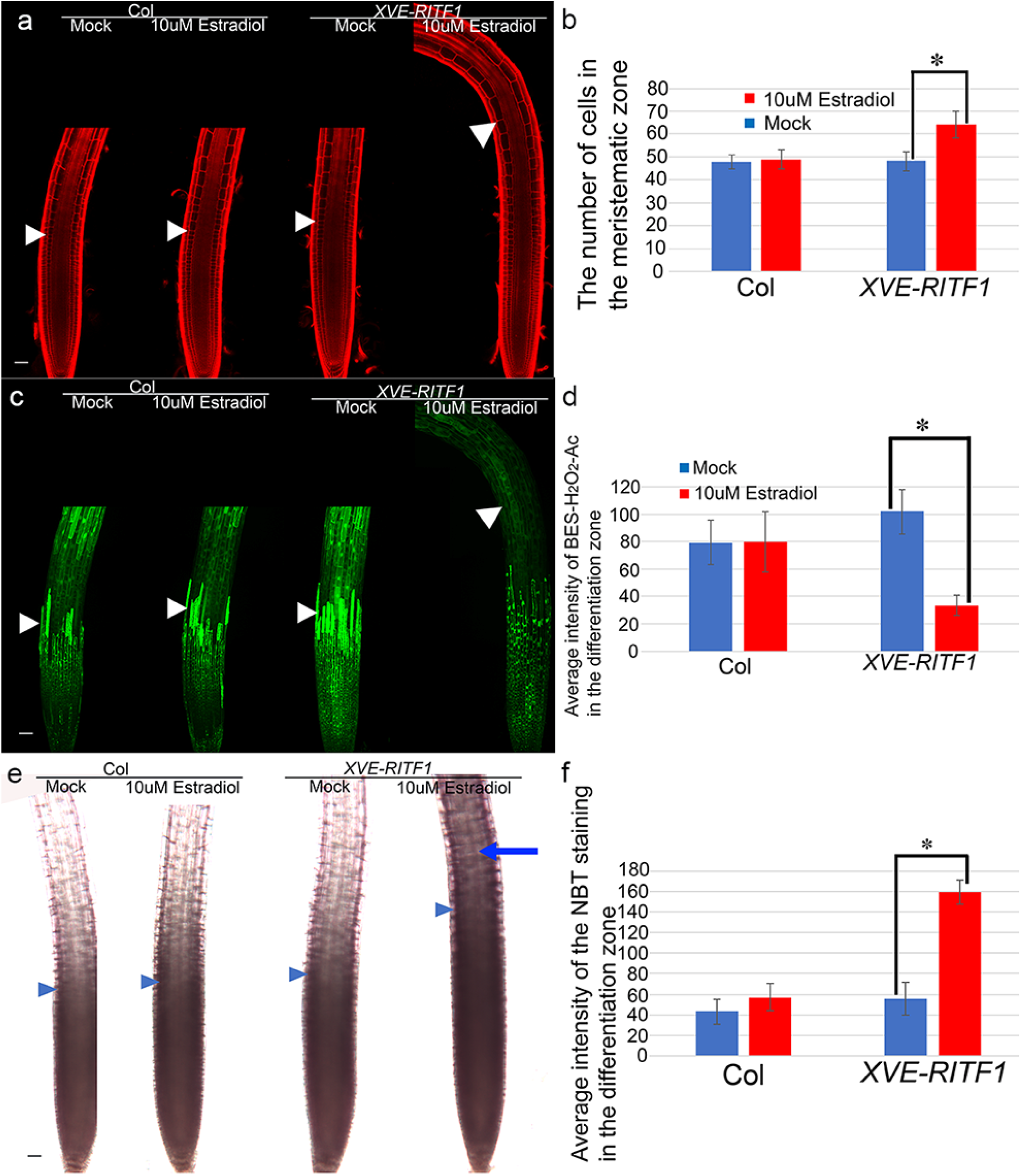
Over-expression of *RITF1*. (a) Confocal images (left WT, right *XVE-RITF1*) of PI stained roots. (b) Number of cells in meristematic zone in wild type (Col) and *XVE-RITF1* 24h after mock or 10µM Estradiol treatment. (N=6, ±SD, and *p<0.001). (c) Confocal images (left *WT, right XVE-RITF1*) of H_2_O_2_-BES-Ac stained roots. (d) Average intensity of BES-H_2_O_2_-Ac in differentiation zone 24h after mock or 10µM Estradiol treatments. (N=6, ±SD, and *p<0.001). (e) Light microscope images of NBT stained roots. Seedlings were grown on MS agar plates for seven days before mock or 10µM estradiol treatment. (f) Average intensity of NBT staining in differentiation zone 24h after mock or 10µM Estradiol treatments. (N≥7, ±SD, and *p<0.001). Scale bar=50µm. White and blue arrow heads show the junction between the meristematic zone and the elongation zone.

Changes in ROS levels could also be detected in the *RITF1* over-expression line (Fig. 4c-4f). H_2_O_2_ levels declined in all three developmental zones upon estradiol treatment (Fig. 4c and 4d). Furthermore, ectopic over-expression of *RITF1* enhanced O_2_^-^ signals in a broader area in the meristematic zone (Fig. 4e) and ectopic O_2_^-^ signals were detected in the elongation and differentiation zone (Fig. 4e, a blue arrow and 4f), where RGF1 receptors are not expressed^6^. This suggests that the *RITF1* gene may control ROS levels without other downstream regulators of the RGF1 signaling pathway.

The larger meristematic zone and alteration of ROS signals by over-expression of *RITF1* strongly suggest that the *RITF1* gene is a primary regulator controlling ROS signaling and meristem size in the RGF1 signaling pathway. Furthermore, the *ritf1* mutant (T-DNA insertion line) has a root growth defect with a smaller meristematic zone (Extended Data Fig. 5c and 5d), and exhibits resistance to RGF1 (Extended Data Fig. 5d and 5e). Taken together, these results indicate that RGF1 modulates ROS levels and root meristem size by controlling expression of the *RITF1* gene.

We have previously reported that UPB1 reduces H_2_O_2_ levels and controls meristem size by down-regulating peroxidase genes in the elongation zone^2^. Our transcriptome analysis didn’t find significant changes in *UPB1* expression upon RGF1 treatment (Extended Data). We did find elevated expression of 5 peroxidase genes (Extended Data), but these are not downstream targets of *UPB1*^2^, suggesting that RGF1 regulates meristem size independently of *UPB1*. However, it is still possible that RGF1 controls meristem size via these peroxidases. To determine if the peroxidase genes upregulated by RGF1 play a role in meristem size control in the RGF1 signaling pathway, we over-expressed two of them (At5g39580 and At4g08780). In neither case did we observe a larger meristematic zone (data not shown).

Identification of the RGF1 peptide and its receptors has provided a novel pathway for regulation of root growth ^3–8^. Initially, it was shown that this signaling pathway regulates root meristem size by enhancing PLT1/2 stability ^3,6,8^. However, the underlying mechanism was not elucidated. We show that RGF1 can modulate ROS levels in the meristematic zone over relatively short time periods and that longer RGF1 treatment results in altered distributions of ROS along the developmental zones of the root. Moreover, PLT2 protein localization correlates with ROS distribution and elevated H_2_O_2_ levels reduce the stability of the PLT2 protein even in the presence of RGF. Finally, we identified a novel transcription factor, *RITF1*, that is induced by RGF1 in the meristematic zone and can alter ROS levels and meristem size. Taken together, our results show that the RGF1 signaling pathway modulates ROS levels and distribution by up-regulating the *RITF1* gene in the meristematic zone.

## Methods

### Plant materials and growth conditions

All *Arabidopsis* mutants and marker lines used in this research are in the Columbia-0 (Col-0) background. The T-DNA insertion line *ritf1* (SALK_081503C) was obtained from the Arabidopsis Biological Resource Center at Ohio State University. The T-DNA insertion was identified at 787 bp downstream of the transcription start site in the ritf1 mutant. Seeds were surface sterilized using 50% (vol/vol) bleach and 0.1% Tween20 (Sigma) for 15 min and then rinsed five times with sterile water. All seeds were plated on standard MS media (1× Murashige and Skoog salt mixture, Caisson Laboratories), 0.5 g/L MES, 1% Sucrose, and 1% Agar (Difco) and adjusted to pH 5.7 with KOH. All plated seeds were stratified at 4°C for 2 d before germination. Seedlings were grown on vertically positioned square plates in a Percival incubator with 16 h of daily illumination at 22 °C.

### Detecting *gPLT2-YFP* and ROS signals

The seedlings of wild type and the *rgfr1/2/3* mutant were grown for six days on MS agar plates, then transferred to MS agar plates containing either water (mock) or 20 nM synthetic sulfated RGF1 peptide (Invitrogen). After RGF1 treatment, seedlings were stained for 2 min in a solution of 200µM NBT in 20mM phosphate buffer (pH 6.1) in the dark and rinsed twice with distilled water. For hydrogen peroxide detection with BES-H_2_O_2_-Ac^16^, seedlings were incubated in 50μM of BES-H_2_O_2_-Ac (WAKO) for 30 min in the dark, then were mounted in 10 mg/mL propidium iodide (PI) in water^2^. Roots were observed using a 20× objective lens under a Zeiss LSM 880 laser scanning confocal microscope. Excitation and detection windows were set as follows: BES-H_2_O_2_-Ac, excitation at 488nm and detection at 500-550 nm; PI staining, excitation at 561 nm and detection at 570-650 nm. Confocal images were processed, stitched, and analyzed using the Fiji package of ImageJ^17^. The maximum projection image was produced from the z-section images of BES-H_2_O_2_-Ac staining. The average intensity of BES-H_2_O_2_-Ac in the meristematic zone was measured in 5 or 6 roots with three biological replicates. Images for NBT staining were obtained using a 10x objective lens under a Leica DM5000-B light microscope. The total intensities of NBT staining in the meristematic zone were measured in 10 roots with three biological replicates using the Fiji software package^17^.

For shorter time course experiments, seedlings of *gPLT2-YFP^11^* were grown on MS agar plates for 6 days, then transferred to MS agar plates containing either water (mock) or 100nM RGF1 peptide. At 6, 8, and 10h after transfer to RGF1 plates, images were taken with a confocal or light microscope after PI, NBT, and BES-H_2_O_2_-Ac staining, as previously described.

### Total RNA preparation, RNA amplification and library preparation for RNA-seq

The *HYP2-GFP^10^* lines were grown on MS plates for 6 days. *HYP2-GFP* seedlings were then transferred into liquid MS media and treated with water (mock) or 100nM RGF1 peptide in 6-well-plates for 1h. After 1h treatment with mock or RGF1, the seedlings were taken out of liquid MS media and transferred onto a 2% agarose plate. Using an ophthalmic scalpel (Feather), the meristematic zone of the seedlings was precisely dissected based on *HYP2-GFP* fluorescence as detected under a dissecting microscope (Axio Zoom, Zeiss). Total RNA was extracted from 20 root sections treated with mock or 100nM RGF1 using the RNeasy Micro Kit (Qiagen). For each treatment, three replicates of the RNA extractions were performed. All total RNA samples were treated with DNaseI during RNA extraction. RNA quality was examined using a 2100 Bioanalyzer (Agilent). The RNA Integrity Number (RIN) was over 9.0 in all samples. The concentration of total RNA was measured by a Qubit (Invitrogen) instrument. For each replicate, 50ng total RNA was amplified using the Ovation RNA-seq System V2 (NuGEN). Following amplification, 3μg of cDNA was fragmented using the Covaris S-Series System. 400ng of the fragmented cDNA with an average size of 400bp was used for library preparation using the Ovation Ultralow System V2 (NuGEN). Illumina sequencing was performed at the Duke Genome Sequencing Shared Resource. The libraries for three biological replicates of mock and RGF1 treated meristematic zones were sequenced on an Illumina HiSeq2000 (100 base-paired reads).

### Differential expression analysis following RGF1 peptide treatment

Illumina sequencing reads were mapped to TAIR10 Arabidopsis genome using Tophat V2.0.7. The parameters used for mapping are as follows : “-N 5 --read-gap-length 5 --read-edit-dist 5 --b2-sensitive -r 100 --mate-std-dev 150 -p 5 -i 5 -I 15000 --min-segment-intron 5 --max-segment-intron 15000 --library-type fr-unstranded”. To select properly mapped reads with unique mapping positions, only alignments with flag of 83, 99, 147 or 163 and a mapping quality score of 50 were kept for further analysis. Mapping positions of these reads were compared with the Araport11 genome annotation (https://www.araport.org/downloads/Araport11_Release_201606/annotation) using HTseq-count generated read count per gene with parameters “--stranded=no --mode=intersection-nonempty”. The raw read counts of miRNA, lncRNA and protein coding genes were then used as input into DESeq2 for differential gene expression analysis. Genes with an adjusted p-value less than or equal to 0.1 were regarded as differentially expressed between RGF treatment and mock. The enriched gene ontology groups among differentially expressed genes were identified as Differences using agriGO (downloaded from http://geneontology.org). A customized GO annotation was used that required a significance level of 0.01 and a minimum mapping entry of 10.

### Plasmid Construction

The coding sequence of the *RITF1* gene (AT2G12646) was amplified using the Phusion High-Fidelity DNA polymerase (New England Biolabs) from a wildtype cDNA library, and then sub-cloned into the *pENTR/D/TOPO* vector (Invitrogen). The following primers were used for *RITF1* amplification: 5’-CACCATGGGAATTCAGAAACCGG-3’ and 5’-TTAACAGAGAGGAGATCGTTG-3’. The sequence of the *RITF1* gene in ***pENTR/D/TOPO*** vector was confirmed by Sanger sequencing. The clone was recombined with the ***pMDC7*** vector^15^ using LR clonase II (Invitrogen) to fuse the estradiol inducible promoter (XVE)^18^ and the coding region of the *RITF1* gene.

## Measurement of meristem size and detection of ROS signals after over-expression of the *RITF1* gene

The ***XVE-RITF1*** construct was transformed into wild type (Col). To measure meristem size and detect ROS signals, two independent lines of *XVE-RITF1* and wild-type were grown on MS media for six days, then transferred to MS media containing DMSO (Mock) and 10μM β-estradiol (Sigma). After 24h treatment with mock or estradiol, meristem size and ROS signals were measured and detected in wild type and the *XVE-RITF1* lines, as described in the previous section.

**Supplementary Information** is available in the online version of the paper.

## Acknowledgments

We thank Jazz Dickinson, Edith Pierre-Jerome, Kevin Lehner, and Cara Winter for comments on the manuscript; Carmen Wilson for help with generating overexpression lines; Keiko Sugimoto for *HYP2-GFP* seed; Yoshikatsu Matsubayashi for the triple mutants of *rgf1/2/3* and *rgfr1/2/3*; Renze Heidstra for *gPLT2-YFP* and *pPLT2-CFP* seed; Nam-Hai Chua for pMDC7 vector; the Duke Genome Sequencing Center for sequencing the Illumina libraries. This work was funded by the Howard Hughes Medical Institute and the Gordon and Betty Moore Foundation (through Grant GBMF3405) and from the NIH (R01-GM043778).

## Author contributions

M.Y. and P.N.B. conceptualized the study; M.Y. performed all experiments; X.H. performed the computational analyses; all authors wrote the paper.

## Author information

Reprints and permissions information is available at www.nature.com/reprint. The authors decleare competing financial interests: details are available in the online version of the paper. Readers are welcome to comment on the online version of the paper. Correspondence and requests for materials should be addressed to P.N.B.

